# Functional partitioning of local and distal gene expression regulation in multiple human tissues

**DOI:** 10.1101/046383

**Authors:** Xuanyao Liu, Hilary K. Finucane, Alexander Gusev, Gaurav Bhatia, Steven Gazal, Luke O’Connor, Brendan Bulik-Sullivan, Fred A. Wright, Patrick F. Sullivan, Benjamin M. Neale, Alkes L. Price

## Abstract

Studies of the genetics of gene expression have served as a key tool for linking genetic variants to phenotypes. Large-scale eQTL mapping studies have identified a large number of local eQTLs, but the molecular mechanism of how genetic variants regulate expression is still unclear, particularly for distal eQTLs, which these studies are not well-powered to detect. In this study, we use a heritability partitioning approach to dissect the functional components of gene regulation. We make use of an existing method, stratified LD score regression, that leverages all variants (not just those that pass stringent significance thresholds) to partition heritability across functional categories, and we extend this method to partition local and distal gene expression heritability in 15 human tissues. The top enriched functional categories in local regulation of peripheral blood gene expression included super enhancers (5.18x), coding regions (3.73x), conserved regions (2.33x) and four histone marks (p<3x10^-7^ for all enrichments); local enrichments were similar across the 15 tissues. We also observed substantial enrichments for distal regulation of peripheral blood gene expression: super enhancers (1.91x), coding regions (4.47x), conserved regions (4.51x) and two histone marks (p<3x10^-7^ for all enrichments). Analyses of the genetic correlation of gene expression across tissues showed that local gene expression regulation is largely shared across tissues, but distal gene expression regulation is highly tissue-specific. Our results elucidate the functional components of the genetic architecture of local and distal gene expression regulation.

## Introduction

Our understanding of functional elements of the human genome has benefitted greatly from the explosion of functional data generated by the ENCODE project and the Roadmap Epigenomics consortium^1,2^. In particular, researchers have gained new insights on the functional effects of genetic variants on many complex diseases and traits^3-11^. In parallel, large-scale eQTL mapping studies have been carried out to catalog genetic variants that affect gene expression in multiple human tissues^12-18^ (reviewed in ref. 19). Gene expression serves as an important intermediate cellular phenotype that affects complex diseases and traits^20-23^, and the functional effects of eQTLs provide another lens through which researchers can investigate molecular mechanisms^9,12-19,24-26^.

However, the functional effects of eQTLs are still largely unclear. On one hand, previous studies have produced different functional characterizations of local eQTLs (Table S1), perhaps due to differences in the sets of annotations analyzed and/or the sample size dependence of approaches that assess enrichment using only top eQTLs. On the other hand, functional characterization of distal eQTLs has been limited^14,15^, since most studies are under-powered to detect distal eQTLs.

In this study, we applied a recently developed method, stratified LD score regression, to partition the heritability of local and distal gene expression regulation across different functional categories^10^. This method makes use of summary association statistics of all genetic variants (not just the top significant variants), and estimates the heritability explained by each functional category while accounting for linkage disequilibrium (LD) to other functional categories; simulations show that this is more powerful than other methods for detecting functional enrichment (Fig. 7 of ref. 10). We extended the method to produce aggregate estimates across all genes for both local and distal gene expression regulation. By applying the method to large gene expression datasets in multiple human tissues, we aimed to comprehensively assess the functional enrichments of genetic variants on local and distal gene expression regulation and shed light on the underlying molecular mechanisms.

## Results

### Heritability enrichment of local gene expression regulation in 15 human tissues

We used an extended version of stratified LD score regression^10^ to partition local gene expression heritability in three data sets spanning 15 human tissues^12,15,18^(Table 1 and Online Methods). We considered 57 functional categories: the 53 baseline categories from ref. 10, and 4 categories based on super enhancers and typical enhancers from ref. 27(see Online Methods). We estimated the enrichment of each functional category, defined as the proportion of heritability in that category divided by the proportion of SNPs in that category. We also report an AUC metric that quantifies how informative each enrichment is; this metric quantifies the fact that larger categories (i.e. spanning a larger fraction of the genome) are more informative than smaller categories at a given enrichment level (see Online Methods).

**Table 1.**
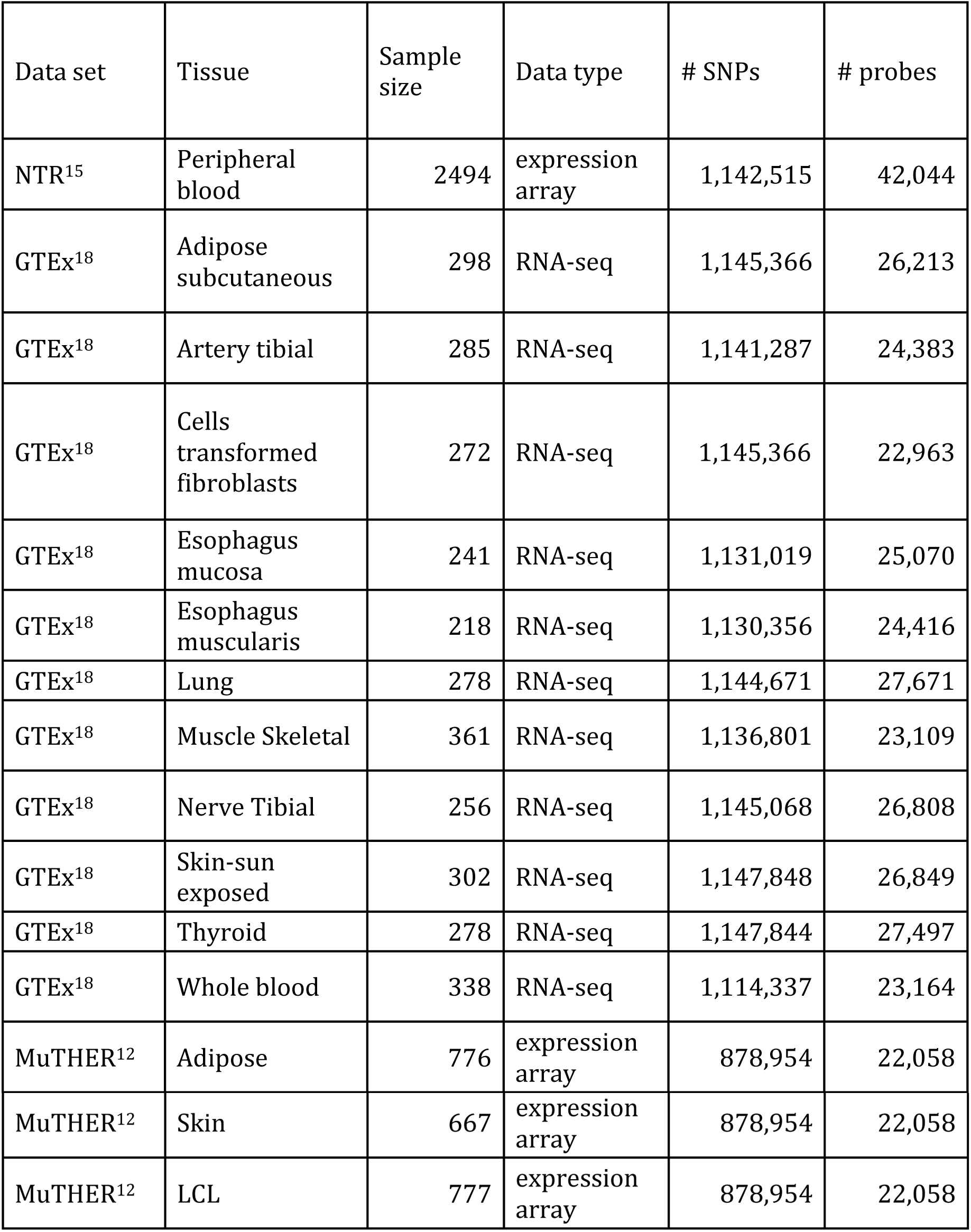
**Gene expression data sets**. We analyzed gene expression data spanning 15 human tissues from 3 data sets. For each tissue we list the sample size, data type, number of SNPs analyzed, and number of probes analyzed. We note that stratified LD score regression restricts to HapMap3 SNPs from the target data set, as a proxy for SNPs with high quality imputation.

We first analyzed the Netherlands Twin Registry (NTR) gene expression array data set, which had the largest sample size (N=2,494) and included only a single tissue type, peripheral blood^15^. Many functional categories were significantly enriched (Figure 1, Table S2). These included several functional enrichments that were not reported in previous studies of gene expression in humans. We observed a large enrichment at Vahedi et al.27 super enhancers (5.18x, p=5.78x10^-7^), supporting the role of super enhancers in gene expression regulation^27,28^. (Hnisz et al.^28^ super enhancers, which span a larger proportion of the genome, produced a smaller but more significant enrichment; Table S2). Conserved regions were also significantly enriched (2.33x, p=9.93x10^-11^). Though the function of conserved regions in gene regulatory programs has previously been reported in yeast^29^, previous evidence of functional enrichments of conserved regions on gene expression in humans is limited: ref. 24 reported a 1.9x enrichment of top eQTLs in conserved elements, but the enrichment was not statistically significant; ref. 25 reported that conserved regions provided “little information” for predicting eQTL. Our results demonstrate that conserved regions indeed play an important role in gene expression regulation.

**Figure 1.**
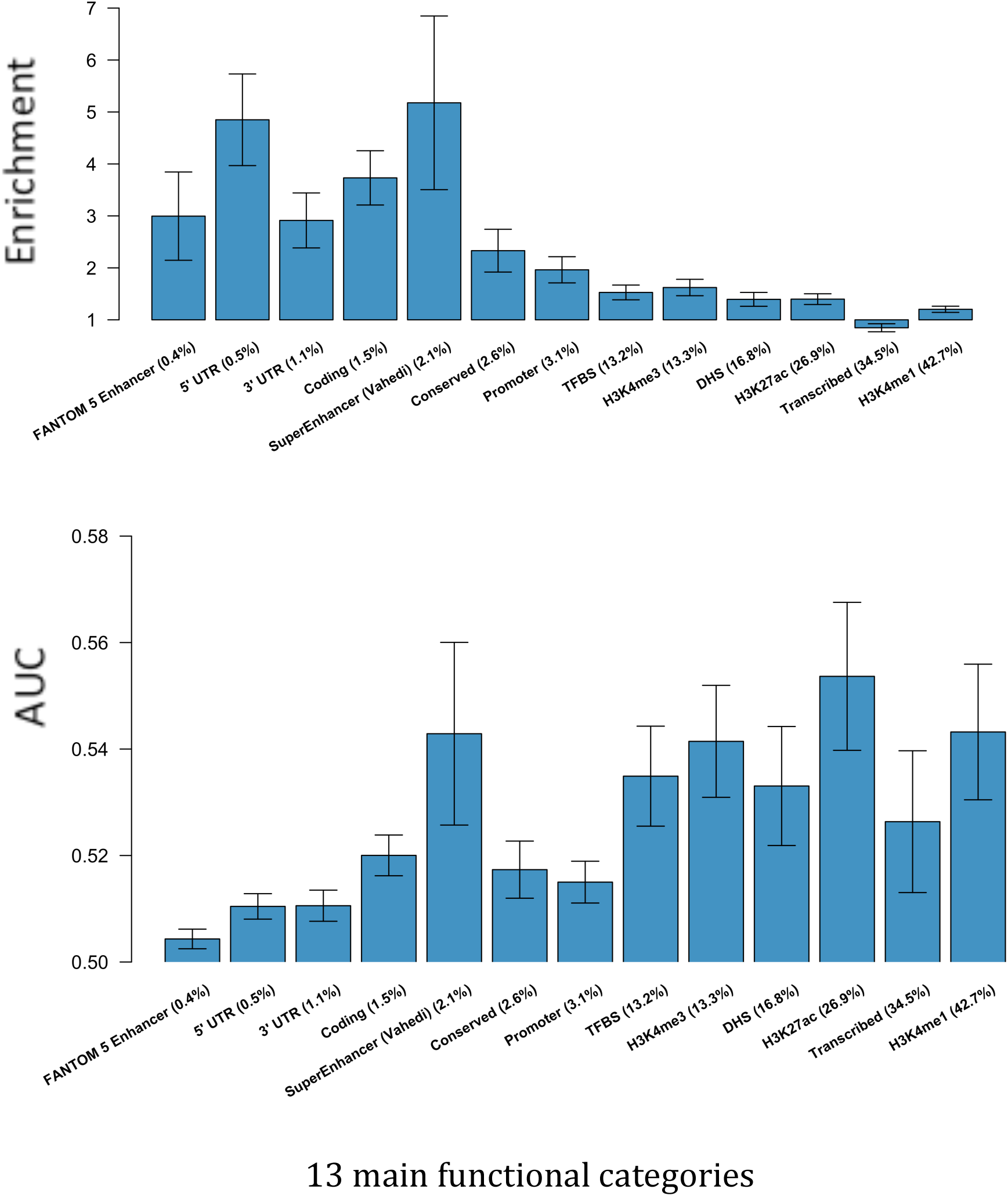
**Functional enrichments of 13 categories for local gene expression regulation in peripheral blood (NTR data set)**. The 13 categories are ordered by the proportion of SNPs in the category. We report (A) The heritability enrichment of each category and (B) the AUC for enrichment of each category. Error bars represent 95% confidence intervals. Numerical results for all 57 categories analyzed are reported in Table S2.

We also confirmed and quantified functional enrichments reported in previous studies of local gene expression regulation in humans. We observed a large enrichment in coding regions (3.73x, p<10^-12^), which confirmed previous findings^9,13^ (Table S1) and is consistent with a recent study reporting that exonic regions are often involved in transcription factor binding^30^. This suggests that the impact of coding variants on complex traits may often be due to changes in expression levels rather than changes in coding sequences. The histone marks H3K4me3, H3K9ac, H3K4me1 and H3K27ac were significantly enriched (enrichment>1.2x, p<3x10^-7^), consistent with previous findings^13,18,25^ (Table S1), confirming the important role of histone marks in local gene expression regulation. Five-prime untranslated regions (5’ UTRs) were also significantly enriched (4.85x, p<10^-12^). This enrichment may be driven by the transcription start site (TSS), which lies inside the 5’ UTR and directly affects transcription, or due to the effect of variants in upstream open reading frames located in the 5’ UTRs on transcript stability^31,32^. We also observed significant enrichments at DNase I hypersensitivity sites (DHS), enhancers, promoters and transcription factor binding sites (TFBS), consistent with previous studies (Table S1).

We extended our analyses to include additional RNA-seq (GTEx) and gene expression array (MuTHER) data sets spanning a total of 15 tissues (Table 1 and Online Methods). The heritability enrichments were highly consistent across the 15 tissues, despite the widely varying sample sizes and different assays (Figure 2, Table S2, Table S3 and Table S4), which indicates that the functional architecture of local gene expression regulation is consistent across different tissues. We note that in contrast to stratified LD score regression, methods for assessing functional enrichment using only top eQTL may be highly sample size dependent (see Discussion).

**Figure 2.**
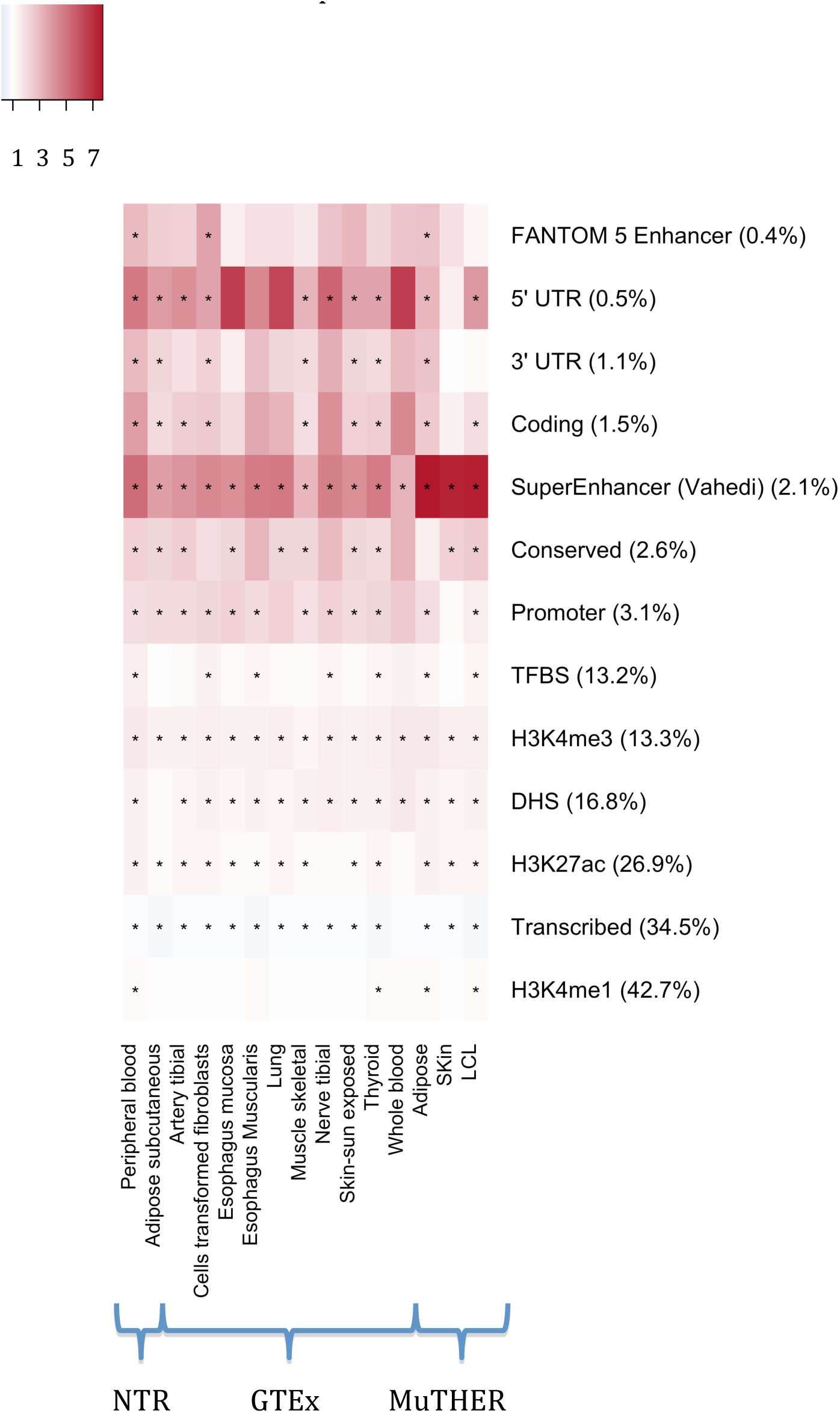
**Functional enrichments of 13 categories for local gene expression regulation in 15 tissues**. Red shading indicates enriched categories (enrichment >1), blue shading indicates depleted categories (enrichment < 1), and * indicates significant enrichment or depletion after correction for 57 hypotheses tested. Numerical results are reported in Table S2, Table S3 and Table S4.

We compared the functional enrichments that we estimated for local gene expression regulation in peripheral blood to functional enrichments that we previously reported for a meta-analysis of 9 independent complex traits^10^ for 53 baseline functional categories. We observed a moderately strong correlation (Pearson r=0.53), as functional categories with larger enrichments for complex traits tend to have larger enrichments for local gene expression regulation (Figure 3). The enrichments in local gene expression regulation tended to be smaller than the enrichments in complex traits: enrichments for 19 (resp. 1) out of 53 categories were significantly smaller (resp. larger) for local gene expression regulation (Table S5). For example, conserved regions were 2.33x enriched in local gene expression regulation vs. 13.31x in complex traits (p=4.36x10^-13^ for difference), coding regions were 3.73x enriched in local gene expression regulation vs. 7.12x in complex traits (p=1.20x10^-4^ for difference), and H3K4me1 were 1.20x enriched in local gene expression regulation vs. 1.86x in complex traits (p=2.29x10^-5^ for difference). Notably, because of the large number of genes in each gene expression dataset, analyzing gene expression as an intermediate phenotype resulted in much smaller standard errors as compared to analyses of complex traits in very large sample sizes, leading to enrichments that were more statistically significant despite their lower magnitude (Figure 3, Table S5). For example, the 1.40x enrichment of H3K27ac in local gene expression was more significant (p<10^-12^) than the 1.82x enrichment of H3K27ac in complex traits (p=4.30x10^-5^). Thus, gene expression data can be a particularly valuable means to assess functionally important genomic regions.

**Figure 3.**
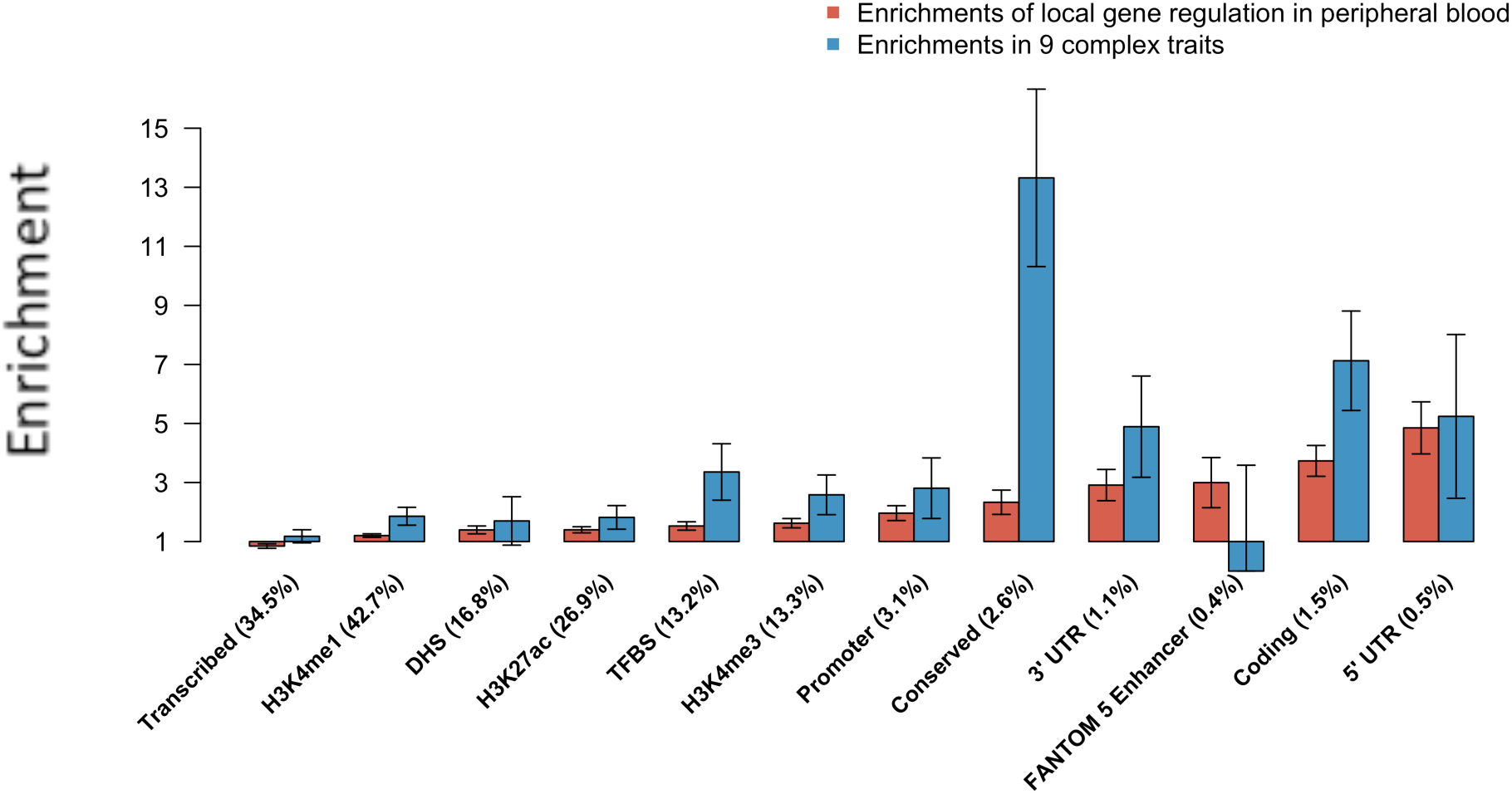
**Comparison of functional enrichments of 13 categories for local gene expression regulation in peripheral blood (NTR data set) vs. 9 complex traits.** Enrichments for local gene expression regulation (red bars) are identical to Figure 1A. Enrichments for complex traits (blue bars) are meta-analyzed enrichments of 9 complex traits and diseases from ref. 10. The categories are ordered by enrichment in the local gene expression analysis. We note that two categories have point estimates with discordant effect directions (enrichment>1 vs. enrichment<1), but in each case the complex trait enrichment is not significantly different from 1. Error bars represent 95% confidence intervals. Numerical results are reported in Table S5.

### Heritability enrichment of distal gene expression regulation in four human tissues

We next used an extended version of stratified LD score regression to partition distal gene expression heritability (see Online Methods). We first analyzed the NTR gene expression array data set^15^. Many functional categories were significantly enriched in the distal analysis (Figure 4, Figure S1 and Table S6). In particular, we again observed significant enrichments at Vahedi et al.^27^ super enhancers (1.91x, p<10^-12^), coding regions (4.47x, p=1.79x10^-11^) and conserved regions (4.51x, p<10^-12^). In addition, two histone marks were significantly enriched: H3K27ac (1.56x, p<10^-12^) and H3K4me3 (1.56x, p=2.29x10^-7^). H3K4me1 and H3K9ac were not significant after correcting for 57 hypotheses tested, but broadly defined H3K4me1 regions (H3K4me1 extended by 500bp; 60.9% of SNPs) explained 98.0% of heritability (1.61x, p<10^-12^). These results highlight the role of histone modifications in distal gene expression regulation. We note that previous results on functional enrichment of distal eQTLs has been limited: ref. 15 reported distal enrichment in 5’ UTR regions (which were nominally enriched in our analyses: 2.96x, p=0.013). Ref. 14 reported enrichment of blood derived distal eQTLs in enhancers regions of myeloid and lymphoid cell lines, and we similarly detected distal enrichment (1.52x, p=8.30x10^-7^) in enhancers as defined by ref. 33. We are not aware of any other previous results on distal enrichment.

**Figure 4.**
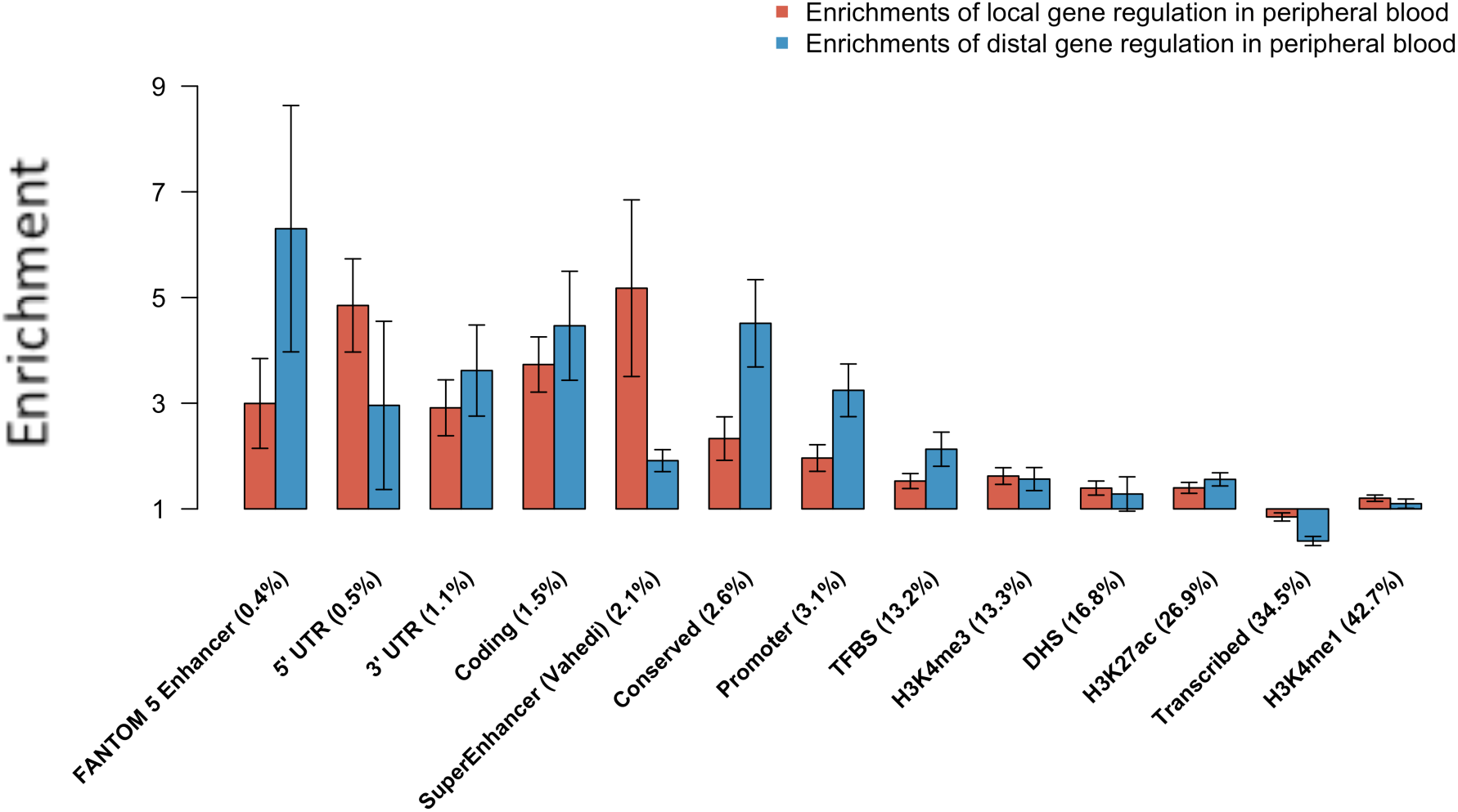
**Comparison of functional enrichments of local and distal gene expression**. We report enrichments for local gene expression regulation (red bars; identical to Figure 1A) and enrichments for distal gene expression (blue bars). The categories are ordered by proportion of SNPs of functional categories. All categories had point estimates with concordant effect directions (enrichment>1 vs. enrichment<1). Error bars represent 95% confidence intervals. Results for the AUC metric are displayed in Figure S1, and numerical results are reported in Table S6.

We compared the distal gene expression regulation enrichments to the local enrichments estimated above, across the 57 categories. We observed a moderately strong correlation (Pearson r=0.48) (Figure 4, Table S6). The enrichments in distal gene expression regulation tended to be somewhat larger than the enrichments in local gene expression regulation; enrichments for 21 (resp. 6) out of 57 categories were significantly larger (resp. smaller) for distal gene expression regulation (Table S6). This suggests that the dearth of previously reported functional enrichments for distal gene expression regulation is due to the low power of approaches based on top distal eQTLs (which most studies are underpowered to detect), and not due to the absence of strong enrichments. Further partitioning of distal regions into intra-chromosomal and inter-chromosomal distal regions produced larger functional enrichments for inter-chromosomal distal regulation than for intra-chromosomal distal regulation (Figure S2 and Table S7). We also analyzed distal enrichment in the MuTHER gene expression array data set (Table 1) and observed many significant enrichments (Figure S3 and Table S8). We did not include the GTEx data set in the distal analysis, due to its smaller sample size.

### Genetic correlation of gene expression between different tissues

We used an extended version of cross-trait LD score regression^34^ to estimate the pairwise genetic correlations of local gene expression between different tissues (see Online Methods). Pairwise genetic correlations were estimated separately in 11 GTEx tissues and in three MuTHER tissues (Figure 5, Table S9 and Table S10). The average pairwise genetic correlation was 0.738. The lowest genetic correlation was observed in whole blood versus esophagus mucosa (r=0.247, s.e.= 0.021). The remaining 57 pairwise correlations were all larger than 0.568, indicating that local regulation of gene expression is highly correlated across tissues, consistent with previous studies^12,18,35-37^. Among the 11 GTEx tissues, whole blood had the lowest genetic correlation with other tissues, with an average genetic correlation of 0.663. This is consistent with GTEx findings that whole blood is an outlier tissue for gene expression patterns and eQTLs discovery^18^.

**Figure 5.**
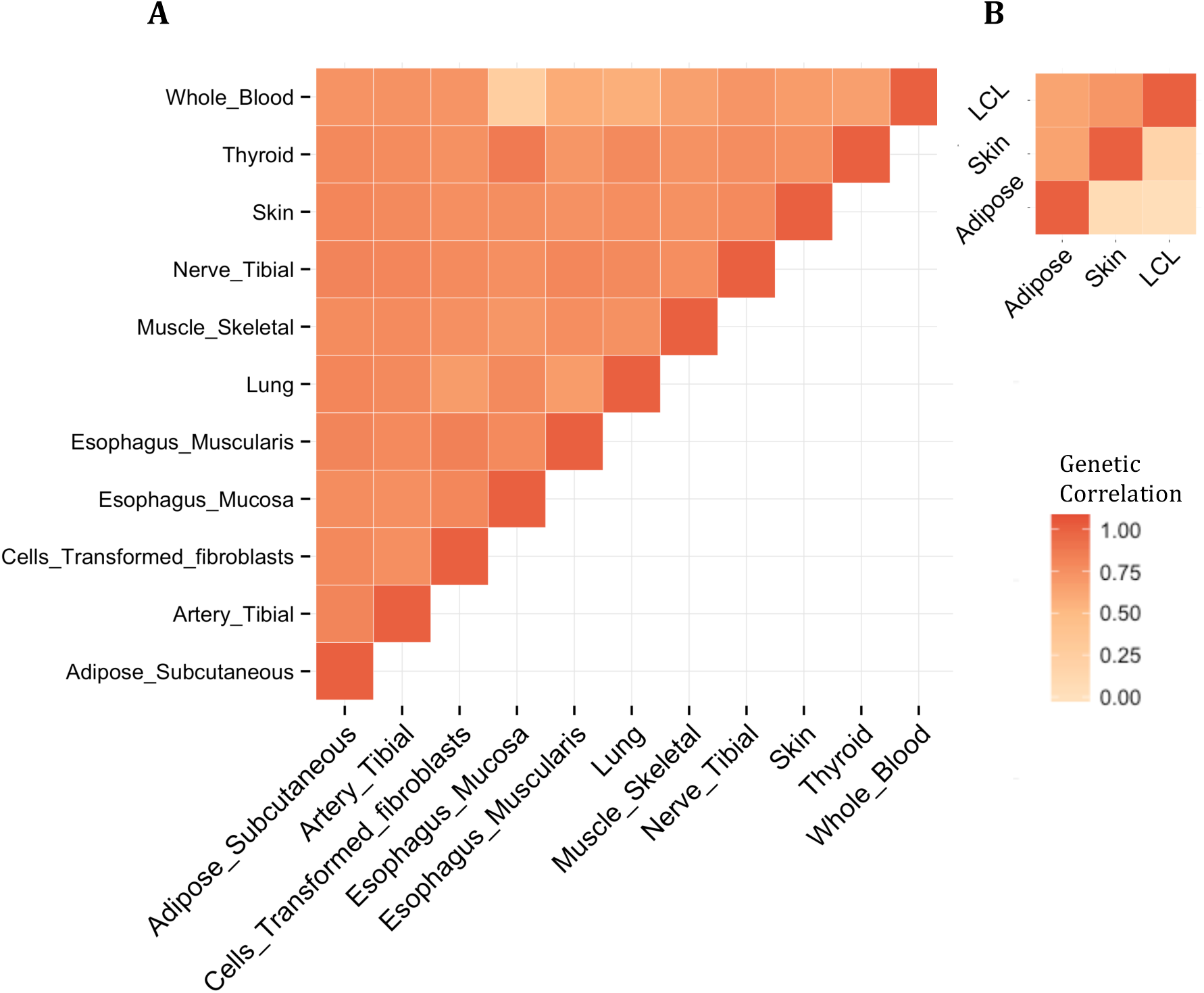
**Pairwise genetic correlation across tissues**. We report (A] Pairwise local genetic correlations across 11 tissues using GTEx data, (B] Local (upper left] and distal (lower right] genetic correlations across 3 tissues using MuTHER data. Numerical results are reported in Table S9 and Table S10.

We also used an extended version of cross-trait LD score regression to estimate the pairwise genetic correlations of distal gene expression in the three MuTHER tissues (Figure 5 and Table S10). The average pairwise genetic correlation was 0.084, indicating that distal regulation of gene expression is highly tissue-specific. This is also consistent with previous work^35^, although few previous studies have investigated the sharing of distal gene expression regulation across tissues due to the low power to detect distal eQTLs.

### Discussion

In this study, we comprehensively investigated functional enrichments for both local and distal gene expression regulation in multiple human tissues by applying an extended version of stratified LD score regression^10^ to large gene expression datasets. We detected widespread functional enrichments for both local and distal gene regulation, including enrichments at super enhancers, coding regions, conserved regions, and several histone marks. We also found that local gene expression regulation is highly genetically correlated across tissues, whereas distal gene expression regulation is highly tissue-specific.

The functional enrichments that we detected for local gene expression regulation were more statistically significant than enrichments that we previously reported for analyses of complex traits in very large sample sizes^10^. This emphasizes the value of studying gene expression as an intermediate phenotype for studying complex diseases and traits, particularly in analyses of functional enrichment. Our systematic investigation of enrichment of local gene expression regulation across 15 tissues identified highly consistent enrichments across tissues, despite the widely varying samples sizes and different assays. This conclusion was possible because the heritability approach employed by stratified LD score regression produces enrichment estimates that are independent of sample size^10^. On the other hand, methods for assessing functional enrichment using only top eQTL may be highly sample size dependent, because the enrichment of associated variants in regulatory annotations may vary with effectsize (see Table 1 of ref. 11). In addition, our results on enrichment of distal gene expression regulation represent a substantial advance over previous results on functional enrichment of distal eQTLs, which were limited by the small number of individually significant distal eQTL detected by previous studies. Our results highlight the advantages of leveraging genome-wide polygenic signals, instead of restricting to top eQTLs, in efforts to identify functional enrichments.

Despite these findings, our work has several limitations. First, stratified LD score regression only models additive effects and cannot capture non-additive effects or epistasis, which may play an important role in gene expression regulation^38-41^. Second, stratified LD score regression is designed for highly polygenic traits and does not take full advantage of non-infinitesimal genetic architectures, which are a particularly likely characteristic of local gene expression regulation. Our highly consistent local enrichments across 15 tissues indicate that the method does produce robust results for local gene expression analyses, but future methods that account for non-infinitesimal genetic architectures might produce even more precise estimates. Third, stratified LD score regression is designed to partition the heritability explained by common variants, but rare variants may also play an important role in gene expression regulation^42^. Fourth, our results on functional enrichment were based on eQTLs and did not consider splicing QTLs (sQTLs), a rich area for future investigation^17,43,44^. Fifth, we detected no significant cell-type-specific local enrichments and only limited cell-type-specific distal enrichments (see Supplementary Note and Figure S4), though similar analyses have detected strong cell-type specific enrichments for complex traits^10^. The absence of local cell-type-specific enrichments is consistent with our observation that local functional enrichments are highly consistent across different tissues, and future analyses may need to restrict to appropriate gene sets (and/or consider sQTLs) to detect cell-type-specific signals. Sixth, we observed smaller enrichments in local gene expression regulation compared to previously reported enrichments in complex traits^10^. Again, future analyses may need to restrict to appropriate gene sets that are relevant to the phenotype of interest (Figure S5). It is also possible that variants in some functional categories contribute to the heritability of complex traits via mechanisms other than local gene expression regulation.

In conclusion, we have identified significantly enriched functional categories for local and distal gene expression regulation in multiple human tissues. These findings shed light on the genetic architecture and molecular mechanism underlying gene expression regulation, and demonstrate that gene expression is an appropriate intermediate phenotype for analyzing functional enrichments of complex diseases and traits.

## URLs

GTEx summary statistics: http://gtexportal.org/home/datasets.

GenABEL and ProbABEL packages: http://www.genable.org.

LD score regression software, including stratified LD score regression and cross-trait LD score regression: http://www.github.com/bulik/ldsc/.

Annotations for 57 functional categories: http://data.broadinstitute.org/alkesgroup/LDSCORE/.

## Online Methods

### Gene expression datasets

We analyzed gene expression in 15 human tissues: peripheral blood from the NTR cohort^15^,11 tissues from GTEx^18^ with sample size larger than 200, and adipose, skin and LCL from the MuTHER cohort^12^ (Table 1). Our analyses required summary association statistics for genome-wide SNPs. For the NTR data set, we used summary statistics from ref. 15, using the t statistics as z scores. For the GTEx data set, we used version 6 of publicly available GTEx summary statistics in local regions (see URLs); for this data set, we considered local regulation only. For the MuTHER data set, we recomputed summary statistics as described in ref. 12. The procedure involves a two-step mixed model-based score test using GenABEL/ProbABEL packages^45,46^ (see URLs). The first step fits a mixed model. The fixed effects include age and batch for adipose and LCL, and age, batch and sample processing for skin. The kinship matrix was built by randomly choosing 10,000 SNPs from the dataset. This step was performed using the *polygenic*() function of the GenABEL software. The second step performs a score test using the ProbABEL software. This step was performed using the -mmscore option of the ProbABEL software.

### Extended version of stratified LD score regression

In a simple linear model,

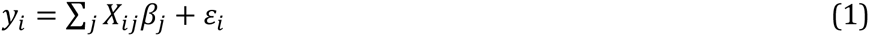

where y**_i_** is a quantitative phenotype in individual *i*, *X_ij_* is the standardized genotype of individual *i* at SNP *j*, *β_j_* is the effectsize of SNP *j*, and *ɛ_i_* is mean-zero noise. The total SNP-heritability is defined as

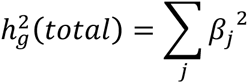

and the SNP-heritability of a category C is defined as

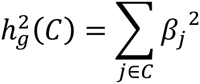

Stratified LD score regression^10^ (see URLs) relies on the fact that LD to a functional category that is enriched for heritability will increase the chi-square association statistics of a SNP more than LD to other categories. More precisely,

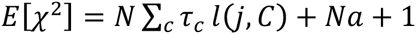

where N is the sample size, *l*(*j*,*C*) is the LD score of SNP j to category C, defined as 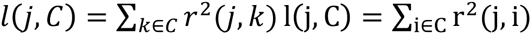, and *a* measures the contribution of confounding biases. (In this study, we employed constrained-intercept LD score regression^34^,in which *a* is fixed at 0.) Performing multiple linear regression of *X*^2^on *l*(*j*,*C*) gives us an estimate 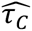 of τ_C_, which represents the per-SNP contribution to heritability of each category C. We estimate 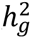(*C*) with

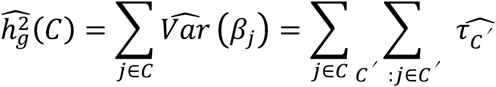

Following ref. 10, we excluded SNPs with chi-square statistics > 80 to reduce variance. We evaluated different chi-square thresholds (excluding SNPs with chi-square > 25, 40, 80 or 300). In both local and distal analyses, the enrichment estimations were not sensitive to the choice of threshold (Table S11 and S12). Following ref.10, we included in our regression only SNPs that appear in HapMap3, which we use as a proxy for SNPs we believe to be confidently imputed. We used the 1000G (phase 1)Europeans^47^ as a reference panel to calculate LD scores. Thus, the LD score *l*(*j*,*C*) for regression SNP *j* is computed using reference SNPs *k* from 1000 Genomes.

We extended stratified LD score regression to analyze gene expression data. For each gene, we applied stratified LD score regression for both local and distal regions of the gene. We defined local regions as the regions within 1Mb of the transcription starting site (TSS) of each gene, and distal regions as the rest of the genome. In local (resp. distal) analyses, both regression SNPs and reference SNPs were restricted to SNPs in local (resp. distal) regions. We also considered a different definition of local regions (within 2Mb of TSS), and determined that the estimates were not sensitive to this choice (Figure S6). We estimated *h_g_*^2^(*C*) and total heritability *h_g_*^2^(*total*) for each gene for both local and distal analyses.

To compute a genome-wide estimate of the proportion of heritability of a category (for either local or distal regions), we sum the category-specific heritability estimates across probes and divide by the sum of total heritability estimates across probes:

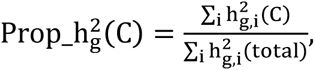

where 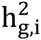(C) denotes the estimated heritability of probe *i* expression in category C, 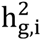(total _!_ denotes the total estimated heritability of probe *i* expression, and the sum is taken over probes i such that 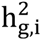(total) > 0. We applied this threshold both because negative heritability is biologically infeasible, and because this reduced estimation noise and resulted in more stable estimates (see Table S13 and Figure S7).

The enrichment of heritability is defined as:

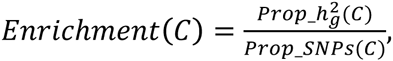

where Prop_SNPs(C) is the proportion of reference SNPs that lie in category C (Table S14).

Standard errors were computed via block-jackknife. In detail, the standard error of Prop_*h_g_*^2^(C) was computed by partitioning the probes by genomic location into 200 blocks and jackknifing on probes. This accounts for possible correlations of nearby probes (analogous to the standard block-jackknife on SNPs employed by stratified LD score regression^10^). The standard error of *Enrichment*(C) was computed by dividing the standard error of *Prop_h_g_*^2^ (*C*) by *Prop_SNPs*(*C*). The statistical significance of enrichment was computed by assuming a normal approximation. We used the significance threshold of 0.05/*n*_c_, where *n*_c_ is the number of categories analyzed, to correct for multiple testing. Open-source software implementing the extended version of stratified LD score regression will be made publicly available as part of the LD score regression software (see URLs) prior to publication

We also computed an AUC metric, which quantifies the fact that larger categories (i.e. spanning a larger fraction of the genome) are more informative than smaller categories at a given enrichment level. In detail, for each category, we calculate the area A under the curve y=f(x), where y is *Prop_h_g_*^2^(*C*), x is *Prop_SNPs*(*C*) (0≤x≤1). The AUC is A, or 1-A if A<0.5 (so that the AUC of a category is equal to the AUC of its complement). The standard error of AUC is calculated as the standard error of Prop_hg2(C) divided by 2.

### Baseline and cell type specific functional categories

The 57 functional categories that we analyzed consist of the 53 basic categories from ref. 10 and an additional 4 categories. The 53 baseline categories from ref. 10 include coding, UTR, promoter and intronic regions, histone marks, DHS, conserved regions, and other annotations, as well as 500bp windows around each of those annotations. The 4 additional categories include super enhancers and typical enhancers in T cells from ref. 27 as well as 500bp windows around each of those annotations. Details of all 57 categories are described in Table S14, and the annotations are publicly available (see URLs).

### Extended version of cross-trait LD score regression

Cross-trait LD score regression^34^ relies on the fact that SNPs with high LD score will a have higher product of z-scores (for two genetically correlated traits) on average than SNPs with low LD scores. More precisely,

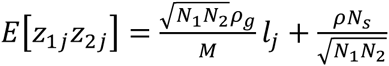

where *N_i_* is the sample sizes for study i, *ρ_g_* is genetic covariance, M is the number of SNPs, l_j_ is the LD score of SNP j, defined as 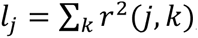, N**_s_** is the number of overlapping samples in the two studies, and ρ is the phenotypic correlation among the common samples.

In the simple model defined by equation (1), let *β_j_* ibe the effect sizes of trait 1 at SNP j and let *γ_j_* be the effect sizes of trait 2 at SNP *j*. The genetic covariance between trait 1 and trait 2 is defined as

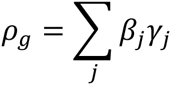

Genetic correlation is defined as

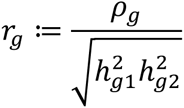

Regressing the product of z scores of two traits on **/***_j_* gives an estimate of *ρ_g_*,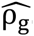. We can also estimate *h_g1_*^2^ and *h_g2_*^2^ from standard LD score regression^48^; the genetic correlation can be estimated from the equation above.

We further extended the method to estimate, for a given pair of tissues, the aggregate genetic correlation of the expression over a large set of common probes. We estimated genetic correlation separately for local and distal regions. For each pair of tissues, for each common probe *i*, we estimated the genetic covariance (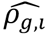), as well as the total heritability of probe *i* in each tissue (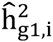 and 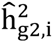). An aggregated genetic correlation across all the shared probes is defined as

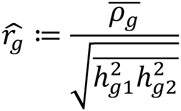

,where 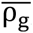, 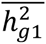 and 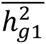 are the averages of 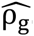, 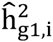 and 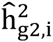 respectively, taken over probes *i* whose 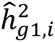 and 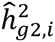 areboth greater than 0. Standard errors of 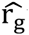 were estimated by dividingthe probes by genomic locations into 200 blocks and performing a block jackknife on the probes, analogous to ref. 34. Open-source software implementing the extended version of cross-trait LD score regression will be made publicly available as part of the LD score regression software (see URLs) prior to publication.

## Acknowledgements

We are grateful to Soumya Raychaudhuri and Xiao Li for helpful discussions. This research was funded by NIH grant R01 MH107649.

## References

1 Kundaje A. et al. Integrative analysis of 111 reference human epigenomes. Nature 518, 317–330 (2015).

2 ENCODE Project Consortium et al. An integrated encyclopedia of DNA elements in the human genome. Nature 489, 57–74 (2012).

3 Ernst J. et al. Mapping and analysis of chromatin state dynamics in nine human cell types. Nature 473, 43–49 (2011).

4 Maurano M. T. et al. Systematic Localization of Common Disease-Associated Variation in Regulatory DNA. 337, 1190–1195 (2012).

5 Trynka G., Sandor C, Han B., Xu H. & Stranger B. E. Chromatin marks identify critical cell types for fine mapping complex trait variants. Genetics (2013).

6 Gusev A. et al. Partitioning Heritability of Regulatory and Cell-Type-Specific Variants across 11 Common Diseases. Am. J. Hum. Genet. 95, 535–552 (2014).

7 Pickrell, J. K. Joint Analysis of Functional Genomic Data and Genome-wide Association Studies of 18 Human Traits. Am.J. Hum. Genet. 94, 559–573 (2014).

8 Kichaev G. et al. Integrating Functional Data to Prioritize Causal Variants in Statistical Fine-Mapping Studies. PLoS Genet 10, e1004722 (2014).

9 Farh, K. K.-H. et al. Genetic and epigenetic fine mapping of causal autoimmune disease variants. Nature 518, 337–343 (2015).

10 Finucane H. K. et al. Partitioning heritability by functional annotation using genome-wide association summary statistics. Nat Genet 47, 1228-1235 (2015).

11 Sveinbjornsson G. et al. Weighting sequence variants based on their annotation increases power of whole-genome association studies. Nat Genet 48, 314–317 (2016).

12 Grundberg E. et al. Mapping cis- and träns-regulatory effects across multiple tissuesintwins. iVatGenet 44, 1084-1089 (2012).

13 Lappalainen T. et al. Transcriptome and genome sequencing uncovers functional variation in humans. Nature 501, 506–511 (2013).

14 Westra H.-J. et al. Systematic identification of träns eQTLs as putative drivers of known disease associations. Nat Genet 45, 1238–1243 (2013).

15 Wright F. A. et al. Heritability and genomics of gene expression in peripheral blood. Nat Genet 46, 430-437 (2014).

16 Battle A. et al. Characterizing the genetic basis of transcriptome diversity through RNA-sequencing of 922 individuals. Genome Research 24, 14–24 (2014).

17 Zhang X. et al. Identification of common genetic variants controlling transcript isoform variation in human whole blood. Nat Genet 47, 345–352 (2015).

18 GTEx Consortium et al. The Genotype-Tissue Expression (GTEx) pilot analysis: Multitissue gene regulation in humans. Science 348, 648–660 (2015).

19 Albert F. W. & Kruglyak L. The role of regulatory variation in complex traits and disease. Nat Rev Genet 16, 197–212 (2015).

20 Davis L. K. et al. Partitioning the Heritability of Tourette Syndrome and Obsessive Compulsive Disorder Reveals Differences in Genetic Architecture. PLoS Genet 9, e1003864 (2013).

21 Torres J. M. et al. Cross-Tissue and Tissue-Specific eQTLs: Partitioning the Heritability of a Complex Trait. Am. J. Hum. Genet. 95, 521–534 (2014).

22 Gamazon E. R. et al. A gene-based association method for mapping traits using reference transcriptome data. Nat Genet 47, 1091-1098 (2015).

23 Gusev A. et al. Integrative approaches for large-scale transcriptome-wide association studies. Nat Genet 48, 245–252 (2016).

24 Veyrieras J.-B. et al. High-Resolution Mapping of Expression-QTLs Yields Insight into Human Gene Regulation. PLoS Genet 4, e1000214-15 (2008).

25 Gaffney D. J. et al. Dissecting the regulatory architecture of gene expression QTLs. Genome Biol 13, R7 (2012).

26 Battle A. et al. Impact of regulatory variation from RNA to protein. Science 347, 664–667 (2015).

27 Vahedi G. et al. Super-enhancers delineate disease-associated regulatory nodes in T cells. Nature 520, 558–562 (2015).

28 Hnisz D. et al. Super-Enhancers in the Control of Cell Identity and Disease. Cell 155, 934–947 (2013).

29 Lee S.-I. et al. Learning a Prior on Regulatory Potential from eQTL Data. PLoS Genet 5, e1000358 (2009).

30 Stergachis A. B. et al. Exonic Transcription Factor Binding Directs Codon Choice and Affects Protein Evolution. Science 342, 1367–1372 (2013).

31 Kervestin S. & Jacobson A. NMD: a multifaceted response to premature translational termination. Nature Reviews Molecular Cell Biology 13, 700–712 (2012).

32 Cenik C. et al. Integrative analysis of RNA, translation, and protein levels reveals distinct regulatory variation across humans. Genome Research 25, 1610-1621 (2015).

33 Hoffman M. M. et al. Integrative annotation of chromatin elements from ENCODE data. Nucl. Acids Res. 41, gks1284-841 (2012).

34 Bulik-Sullivan, B. et al. An atlas of genetic correlations across human diseases and traits. Nat Genet 47, 1236–1241 (2015).

35 Price A. L. et al. Single-Tissue and Cross-Tissue Heritability of Gene Expression Via Identity-by-Descent in Related or Unrelated Individuals. PLoS Genet 7, e1001317-9 (2011).

36 Nica A. C. et al. The Architecture of Gene Regulatory Variation across Multiple Human Tissues: The MuTHER Study. PLoS Genet 7, e1002003 (2011).

37 Flutre T., Wen X., Pritchard J. & Stephens M. A Statistical Framework for Joint eQTL Analysis in Multiple Tissues. PLoS Genet 9, e1003486 (2013).

38 Lappalainen T., Montgomery S. B., Nica A. C. & Dermitzakis E. T. Epistatic Selection between Coding and Regulatory Variation in Human Evolution and Disease. Am. J. Hum. Genet. 89, 459–463 (2011).

39 Hemani G. et al. Detection and replication of epistasis influencing transcription in humans. Nature 508, 249–253 (2014).

40 Wood A. R. et al. Another explanation for apparent epistasis. Nature 514, E3-E5 (2014).

41 Buil A. et al. Gene-gene and gene-environment interactions detected by transcriptome sequence analysis in twins. Nat Genet 47, 88–91 (2015).

42 Zhao J. et al. A Burden of Rare Variants Associated with Extremes of Gene Expression in Human Peripheral Blood. Am. J. Hum. Genet. 98, 299–309 (2016).

43 Ongen H. & Dermitzakis E. T. Alternative Splicing QTLs in European and African Populations. Am. J. Hum. Genet. 97, 567–575 (2015).

44 Gutierrez-Arcelus, M. et al. Tissue-Specific Effects of Genetic and Epigenetic Variation on Gene Regulation and Splicing. PLoS Genet 11, e1004958 (2015).

45 Aulchenko Y. S., de Koning, D.-J. & Haley C. Genomewide Rapid Association Using Mixed Model and Regression: A Fast and Simple Method For Genomewide Pedigree-Based Quantitative Trait Loci Association Analysis. Genetics 177, 577–585 (2007).

46 Chen, W.-M. & Abecasis G. R. Family-Based Association Tests for Genomewide Association Scans. Am. J. Hum. Genet. 81, 913–926 (2007).

47 1000 Genomes Project Consortium et al. An integrated map of genetic variation from 1,092 human genomes. Nature 491, 56–65 (2012).

48 Bulik-Sullivan, B. K. et al. LD Score regression distinguishes confounding from polygenicity in genome-wide association studies. Nat Genet 47, 291–295 (2015).

